# WheatNet: A genome-scale functional network for hexaploid bread wheat, *Triticum aestivum*

**DOI:** 10.1101/105098

**Authors:** Tak Lee, Sohyun Hwang, Chan Yeong Kim, Hongseok Shim, Hyojin Kim, Pamela C. Ronald, Edward M. Marcotte, Insuk Lee

## Abstract

Gene networks provide a system-level overview of genetic organizations and enable the dissection of functional modules underlying complex traits. Here we report the generation of WheatNet, the first genome-scale functional network for T. aestivum and a companion web server (www.inetbio.org/wheatnet). WheatNet was constructed by integrating 20 distinct genomics datasets, including 156,000 wheat-specific co-expression links mined from 1,929 microarray data. A unique feature of WheatNet is that each network node represents either a single gene or a group of genes. We computationally partitioned gene groups mimicking homeologous genes by clustering 99,386 wheat genes, resulting in 20,248 gene groups comprising 63,401 genes and 35,985 individual genes. Thus, WheatNet was constructed using 56,233 nodes, and the final integrated network has 20,230 nodes and 567,000 edges. The edge information of the integrated WheatNet and all 20 component networks are available for download.

## Dear Editor

Gene networks provide a system-level overview of genetic organizations and enable the dissection of functional modules underlying complex traits. With diverse genomics data, a genome-scale network, which facilitates the prediction of novel candidate genes for a trait, can be constructed. Network-based predictions have been useful in the model plant *Arabidopsis thaliana* (Lee et al., 2010). However, such a predictive gene network is not yet available for bread wheat, *Triticum aestivum*, an important staple food crop accounting for approximately 20% of the world’s daily food consumption. Bread wheat also serves as a model for studying polyploidy in plants.

Some of the reasons that functional genomics studies on bread wheat have lagged behind those on other crops include the large genome of bread wheat (~17 Gb) and its polyploidy nature, which complicates genetic analysis. However, recent advances in wheat research have considerably improved genome assembly and gene models (International Wheat Genome Sequencing, 2014). Furthermore, the discovery and application of genome editing (Upadhyay et al., 2013) and TILLING technologies (Uauy et al., 2009) have enabled targeted knockout in wheat protoplasts and whole plants. These developments have set the stage for the application of reverse genetics approaches for the functional characterization of wheat genes.

Here we present WheatNet, the first genome-scale functional gene network for *T. aestivum* and a companion web server (www.inetbio.org/wheatnet), which provides network information and generates network-based functional hypotheses. WheatNet was constructed by integrating 20 distinct genomics datasets (**Supplemental Table 1**), including 156,000 wheat-specific co-expression links mined from 1,929 DNA microarray datasets (**Supplemental Table 2**). A unique feature of WheatNet is that each network node represents either a single gene or a group of genes. An allopolyploid wheat genome contains three homeologous chromosome sets―A, B, and D―that originate from three closely related species *Triticum urartu*, *Aegilops speltoides*, and *Aegilops tauschii*, respectively (International Wheat Genome Sequencing, 2014). Therefore, the wheat genome contains many homologous genes between the three ancestral chromosome sets. Because homeologs are likely to have redundant functions, collapsing homeologs into a single network node would facilitate the network analysis by reducing network complexity. Unfortunately, comprehensive definitions of wheat homeologous relationships are not yet available. Therefore, we computationally partitioned “gene groups” mimicking homeologous genes by clustering 99,386 wheat genes, resulting in 20,248 gene groups comprising 63,401 genes, and 35,985 individual genes. WheatNet was thus constructed using 56,233 nodes; the final network has 20,230 nodes and 567,000 edges, integrating 20 sources of functional evidence linking pairs of genes (**Supplemental Methods**). The edge information of the integrated WheatNet and all 20 component networks are available for download.

To assess WheatNet, we used biological process annotations by agriGO (Du et al., 2010), which are moderately distinct from the dataset used for network training (~38% gene pairs by shared agriGO annotations overlap the training data) and one of the few other large-scale wheat annotation sets available for testing. To help reduce bias, we excluded agriGO terms that annotate more than 300 wheat genes. Next, the accuracy of functional gene pairs by WheatNet or by random chance was measured using the proportion of gene pairs that share agriGO annotations for different coverage of the coding genome. We observed strong performance by WheatNet, in which a network covering approximately 20% of all genes map functional gene pairs with about 40% accuracy (**Supplemental Figure 1**). The quality of WheatNet was further evaluated by the degree of connectivity among genes involved in a particular biological process. Considering that genes for the same complex traits are more likely to be functionally coupled, high connectivity among known genes for a trait would support the quality of functional networks. We tested network connectivity for a group of genes based on two measures: (i) the number of edges among gene members (i.e., within-group edge count) and (ii) the number of network neighbors that overlap among group members (i.e., network neighbor overlap). We used genes for two complex traits derived from proteomics studies: 45 genes with differential protein expression after *Blumeria graminis* f. sp. *tritici* infection (Mandal et al., 2014) and 17 genes with differential protein expression under drought conditions (Cheng et al., 2015). The significance of network connectivity was also measured based on a null distribution from 1000 random gene sets of the same size. We found that the connectivity among each trait’s genes was significantly higher than by random chance (**Figure 1A–B**). We consistently observed network communities of genes for both traits (**Figure 1C–D**). We conclude that WheatNet successfully predicts additional genes that are involved in a given trait.

**Figure 1.**
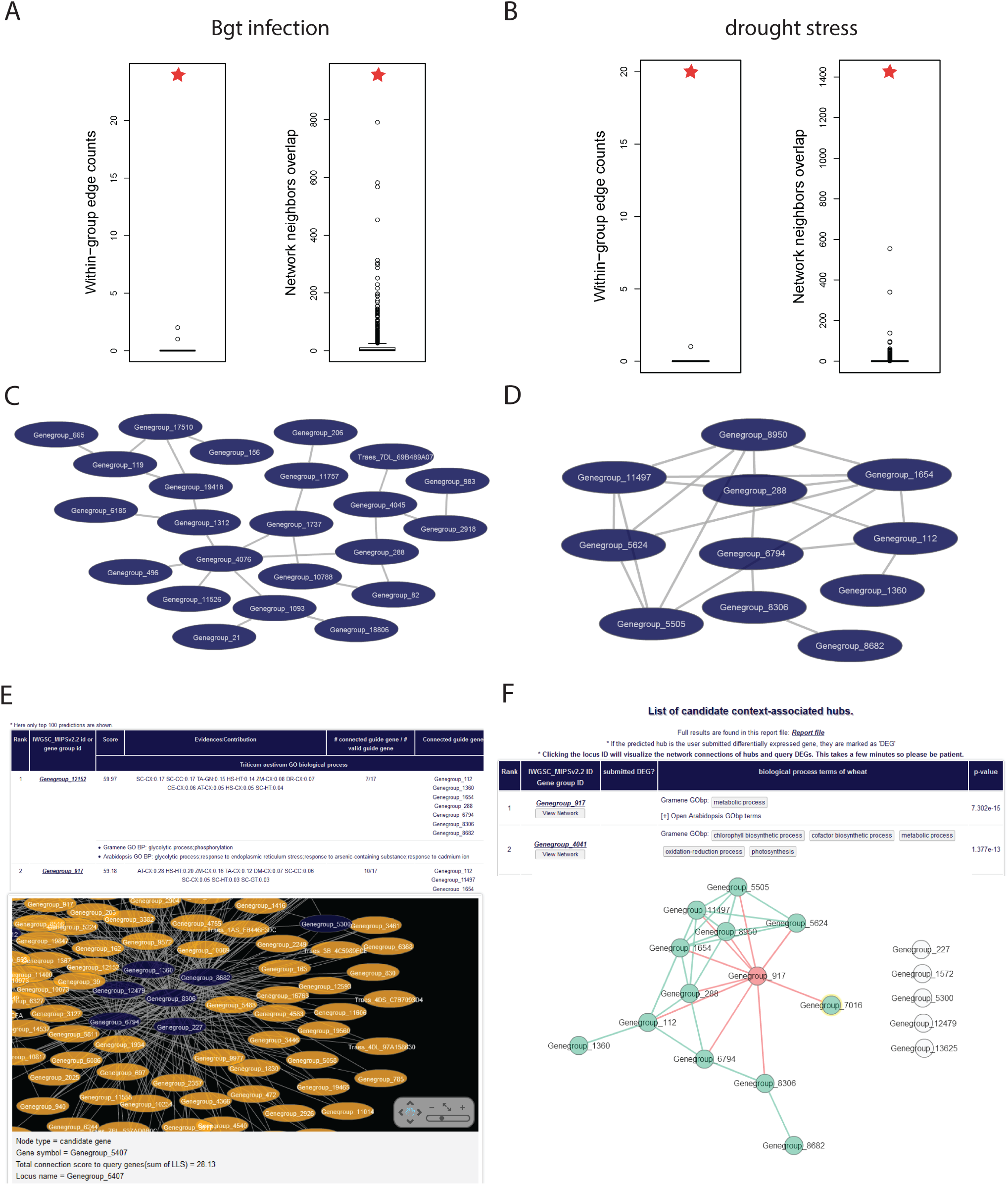
**Overview of WheatNet** Degree of connectivity (A) among 45 genes for response to Blumeria graminis f. sp. Tritici infection and (B) among 17 genes for response to drought stress were measured by determining the number of edges among group members (i.e., within-group edge count) or the number of network neighbors that overlapped among group members (i.e., network neighbor overlap) by using WheatNet (red stars) or 1000 random gene sets having the same size (black circles). Largest components of networks of (C) the genes for response to B. graminis f. sp. Tritici infection and (D) those for response to drought stress by WheatNet. (E) Results of gene prioritization by direct neighborhood method. The top 100 candidate genes and associated Gene Ontology terms are listed in a table. In addition, the network of guide genes and candidate genes is shown. (F) The results of gene prioritization by the context-associated hub method. The top 100 predictions and associated Gene Ontology terms are listed in a table. By clicking each candidate gene, users can view a network composed of the hub gene and connected differentially expressed genes.

The WheatNet web server provides two options for prioritizing genes for wheat traits: (i) direct neighbors in the gene network and (ii) context-associated hubs (CAHs). In the first approach, a user submits genes known for a trait that can guide network searches for new candidate genes. New genes are then ranked by the strength of evidence connecting them to the “guide genes,” measured for each candidate gene as the sum of network edge scores from that gene to the guide genes. The result page provides the ranked list of candidates and a visualization of the local guide gene network using the Cytoscape web tool (Lopes et al., 2010) (**Figure 1E**). To provide functional clues for candidate genes, WheatNet provides available wheat and *Arabidopsis* gene annotations from the Gene Ontology biological process (GOBP) (**Supplemental Methods**).

In the second approach, users exploit gene expression data related to a trait of interest. Gene expression profiles are one of the most common types of genomic data, and differential expression analysis provides many genes that are potentially associated with given traits such as abiotic and biotic stresses. However, many genes that are associated with stress conditions are not differentially expressed. Hypothesizing that a gene associated with many differentially expressed genes (DEGs) in stress (i.e., CAHs) is likely to be responsible for responses to the given stress condition, we prioritized genes by connections to the context-associated DEGs. To conduct CAH prioritization, we first defined a subnetwork that comprises a hub gene and all of its network neighbors in WheatNet. For the gene prioritization, we considered only subnetworks with hub genes that have at least 50 network neighbors. Assuming that DEGs are representative genes for a relevant biological context, we prioritized hub genes based on the enrichment of their network neighbors for the DEGs, measured using Fisher’s exact test. The hub genes with significant enrichment (*P* < 0.01) of network neighbors for the DEGs are considered as CAHs and are presented as candidate genes for the context-associated trait. Similar to the network direct neighborhood search, all candidate genes are appended by GOBP annotations for wheat genes and for Arabidopsis orthologs. In addition, users can access a network view of a CAH and its connected DEGs by clicking each candidate gene (**Figure 1F**).

The WheatNet predictions by each of the network-based gene prioritization methods were validated as follows: For the network direct neighborhood method, we evaluated the new candidate genes for drought stress response that were predicted by submitting 17 genes with differential protein expression under drought conditions (Cheng et al., 2015) as guide genes.

We hypothesized that novel candidate genes for drought response are also likely to be expressed differentially under drought conditions. Thus, we investigated the enrichment of candidate drought response genes from DEGs under drought conditions. We generated a set of 2,346 DEGs under drought condition based on genes that showed more than 4-fold changes in expression levels at *P* < 0.01 (SRP045409 of NCBI Sequence Read Archive) (Liu et al., 2015). We found 15 drought-condition DEGs among the top 50 candidate genes by the network direct neighborhood method, which indicates more than 7-fold enrichment over predictions by random chance (15/50 = 0.3 by WheatNet vs. 2346/56233 = 0.042 by random chance). For the CAH method, we evaluated the candidate genes for *Fusarium graminearum* infection response that were predicted by submitting 837 DEGs after infection with *F. graminearum* (GSE54551 of NCBI Gene Expression Omnibus database) (Wojcik et al., 2015) as user input data. We found that the top 100 candidates by CAHs were significantly enriched for GOBP annotations relevant to fungus infection based on Arabidopsis orthologs: ‘response to chitin’ (GO:0010200, *P* = 9.72 × 10^-31^), ‘regulation of plant-type hypersensitive response’ (GO:0010363, *P* = 8.20 × 10^-21^), ‘defense response to fungus’ (GO:0050832, *P* = 1.73 × 10^-20^), ‘response to fungus’ (GO:0009620, *P* = 1.03 × 10^-8^), and ‘detection of biotic stimulus’ (GO:0009595, *P* = 3.43 × 10^-5^).

These results indicate that WheatNet can effectively prioritize novel candidate genes for complex traits, including those governing abiotic and biotic stress responses, by using multiple network-based methods, which can be easily performed by simple submission of input data in the companion web server. WheatNet should be a useful resource of systems biology and predictive genetics for the wheat research community.

## Author contributions

T.L. and S.H. developed the network model and conducted bioinformatics analysis. C.Y.K., H.S., and H.K. assisted data analysis for network modeling. P.C.R., E.M.M., and I.L. designed and supervised the study. T.L. and I.L. drafted the manuscript. P.C.R., and E.M.M., edited the manuscript.

## Acknowledgments

This work was supported by grants from the National Research Foundation of Korea (2012M3A9B4028641, 2012M3A9C7050151, and 2015R1A2A1A15055859) to I.L. This work was supported by a grant to PCR and EMM from NSF (1237975) and from the Welch Foundation (F1515) to EMM. The work conducted by the US Department of Energy Joint Genome Institute was supported by the Office of Science of the US Department of Energy under Contract no. DE-AC02-05CH11231.

We thank Jorge Dubcovsky, Ksenia Krasileva, Kelly Eversole and Catherine Feuillet for helpful discussions.

## Supplemental Methods

### Genes and gene groups for nodes of WheatNet

Genes for WheatNet are based on gene models by the International Wheat Genome Sequencing Consortium MIPS annotation version 2.2. In the allopolyploid genome of Triticum aestivum, three homeologous chromosome sets―A, B, and D―originate from closely related species *Triticum urartu*, a grass relative *Aegilops speltoides*, and *Aegilops tauschii*, respectively (International Wheat Genome Sequencing, 2014). Consequently, the wheat genome harbors many homologous genes between the three ancestral chromosome sets. Since the homeologs are likely to carry redundant functions, clustering a group of homeologs into a single network node would facilitate network analysis by reducing network complexity. Unfortunately, no comprehensive database of wheat homeologous relationships is yet available. Therefore, we computationally established “gene groups” as mimicking the homeologous groups. We hypothesized that, since homeologs arise from interspecies hybridization of chromosomes, they should resemble orthologous relationships between species. Therefore, we used orthoMCL (Li et al., 2003) to search for groups of homologs among the three chromosome sets. We do not expect these gene groups to completely capture the homeologous relationships, but they should serve to approximate the ancestral homology. Of the 99,386 wheat genes, 63,401 were clustered into 20,248 gene groups. WheatNet was thus constructed on a total of 56,233 nodes, representing the 20,248 gene groups plus 35,985 individual genes. (Unless otherwise specified, we use the word “gene” to refer to a network node that might be either a single gene or a gene group.)

### Benchmarking inferred functional linkages between wheat genes

Various types of genomics data were analyzed to infer functional associations between wheat genes by using supervised machine learning, which requires gold-standard reference data for training the model. The positive gold-standard set of functional links between wheat genes was derived from Gramene Gene Ontology biological process (GOBP) terms (release 49) (Tello-Ruiz et al., 2016) by all-versus-all pairing of member genes for each GO term. Because the number of gene pairs for each term exponentially increases with the number of member genes, the resultant set of gold-standard functional gene pairs could be severely biased toward GO terms with many member genes. To avoid this functional bias, GO terms with more than 200 member genes were excluded. We also generated the negative gold-standard data set by pairing genes annotated by different GO terms. The resultant positive and negative gold-standard data sets comprise 434,368 and 73,273,643 gene pairs, respectively, among 12,142 wheat genes.

Functional linkages between wheat genes inferred from various genomics data were evaluated by using a log likelihood scoring scheme (LLS) (Lee et al., 2004). The LLS is calculated as follows:

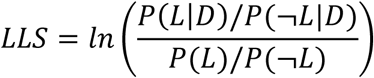

Where P(L|D) and P(¬L|D) are the probability of the positive and negative gold-standard linkages for a given biological data, respectively. In a sense, LLS measures the enrichment of the positive gold-standard links among ones inferred from the given biological data compared to those obtained by random chance. If LLS is above zero, the given pair of genes is likely to operate for the same biological process.

### Integrating the functional linkages from different data sources

To construct the gene network, we integrated functional gene-gene links inferred from 20 different genomics datasets (**Supplementary Table 1**). Since these genomics datasets are not entirely independent from each other, we integrated multiple scores from different datasets using a previously described weighted sum (WS) method (Lehner and Lee, 2008). We found that integrating scores in a weighted fashion generally yields better results than those by *naïve* Bayes integration, in which all scores are summed up with full weights. WS is calculated as follows:

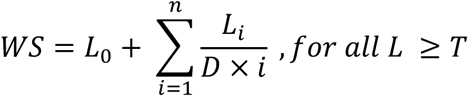

where L_0_ is the best LLS of all available scores for a given functional link. Rank indices from *i* to *n* are assigned to the rest of the LLSs. Next, the scores are summed up after dividing them by the assigned index number and multiplied by a weight factor D if the given LLS is higher than a score threshold T. In this method, weighted sum yields the best LLS with the full weight, and all others with partial weights show decrement along rank order.

### Inferring functional links from co-expression analysis

Genes for related biological processes are likely to have similar expression dynamics across many biological contexts. This co-expression pattern can be exploited to infer functional relationships between genes. We analyzed gene expression data collected using the Affymetrix array platform (GPL3802) and deposited in the Gene Expression Omnibus (GEO) database (Barrett et al., 2013), because it supports the largest number of samples. We converted the downloaded raw data into a matrix of genes (rows) and samples (columns) for each GEO series (GSE), and then normalized the data using MAS5 software. We then calculated the Pearson correlation coefficient (PCC) across these preprocessed expression vectors for each pair of genes in order to measure their co-expression. When analyzing co-expression among genes falling in homeologous gene groups, we retained the highest PCC scores among individual gene pairs. The co-expression links were ranked according to PCC score, and LLS scores were calculated for consecutive bins of 1000 gene pairs. The resulting relationship between PCC and LLS was fit with a regression model, and LLSs assigned to individual gene pairs based on the derived model. GEO series with fewer than 12 samples were excluded from the co-expression analysis owing to the high probability of promiscuous functional links supported by insignificant correlation across low numbers of samples. We inferred functional gene networks from nine GEO series (**Supplemental Table 2**), which were integrated into a component network for WheatNet, TA-CX (see **Supplemental Table 1**). We also analyzed several expression datasets based on high-throughput sequencing, but none showed strong correlation between PCC and LLS, and we did not incorporate these data into the network, although the rapid growth in RNA-sequencing data in the database suggests this will likely become a valuable source of functional evidence in the future.

### Inferring network links from gene neighborhood method by using metagenome assemblies

Functional associations between wheat genes can be inferred by measuring the proximities of their orthologous genes in bacterial genomes, a method that takes advantage of the trend for genes found to be neighbors across many bacterial genomes to reside in the same operon, and thus to generally operate in the same biological process (Overbeek et al., 1999; Shin et al., 2014). Previously, we analyzed gene neighborhoods across completely sequenced bacterial genomes. We adapted the method for metagenomics data spanning hundreds of thousands of bacterial genomes. Metagenome assemblies of 16 body sites of humans were downloaded from the Human Microbiome Project (HMP) (Human Microbiome Project, 2012), and global ocean microbiome assemblies were obtained from TARA Oceans study (Sunagawa et al., 2015). Wheat protein sequences were aligned to the metagenome assemblies by using the BLASTX alignment mode in DIAMOND (Buchfink et al., 2015) with the ‘sensitive’ option. The score of each gene pair (S) was calculated as follows:

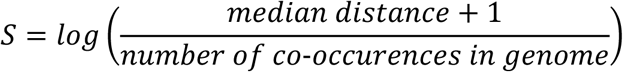

We expected that gene neighborhood evidence across the human symbiotic bacteria and marine bacteria would be complementary, given the many differences in the two environments and taxa, and derived two independent networks, one based on the HMP data and the other based on the TARA Oceans study data. We combined the networks into a single gene neighborhood network (TA-GN in **Supplemental Table 1**) for integration with other datatypes.

### Orthologous functional links transferred from other organisms

Interologs are evolutionarily conserved interactions between proteins whose homologous proteins in other organisms also interact with each other (Walhout et al., 2000). Similarly, associalogs (Kim et al., 2013) are conserved functional associations between genes whose orthologs in other organisms are also functionally coupled. WheatNet component networks based on associalogs were transferred from our previously published networks for seven organisms: YeastNet for *Saccharomyces cerevisiae* (Kim et al., 2014), AraNet for *Arabidopsis thaliana* (Lee et al., 2015c), RiceNet for *Oryza sativa* (Lee et al., 2015b), FlyNet for *Drosophila melanogaster* (Shin et al., 2015), WormNet for *Caenorhabditis elegans* (Cho et al., 2014), DanioNet for *Danio rerio* (Shim et al., 2016), MouseNet for *Mus musculus* (Kim et al., 2016), and HumanNet for *Homo sapiens* (Lee et al., 2011). In addition, associalogs from an unpublished functional network for *Zea mays* were integrated into WheatNet. To transfer associalogs from the above organisms, orthologous wheat proteins were identified using the Inparanoid algorithm, which includes in-paralogs as co-orthologs (O'Brien et al., 2005). We found that associalogs based on wheat in-paralogs of *Z. mays* genes increased the probability of introducing false functional links, probably because of the complexity of the *Z. mays* genome (Lee et al., 2015a). Therefore, we transferred associalogs from *Z. mays* networks by orthology relationships based on the bi-directional best BLASTP hits only, not the in-paralog expansion offered by Inparanoid.

### Functional annotations of wheat genes for the web service

For functional annotation of wheat genes in the web server, we used GOBP terms as assigned by Gramene (release 49) (Tello-Ruiz et al., 2016). *Arabidopsis* GOBP terms were also used to annotate functions of wheat genes based on orthology, as identified using bi-directional best BLASTP hits. We excluded GO annotations whose support derived only from non-traceable author statements with no supporting data.

---

**Supplemental Table 1.**
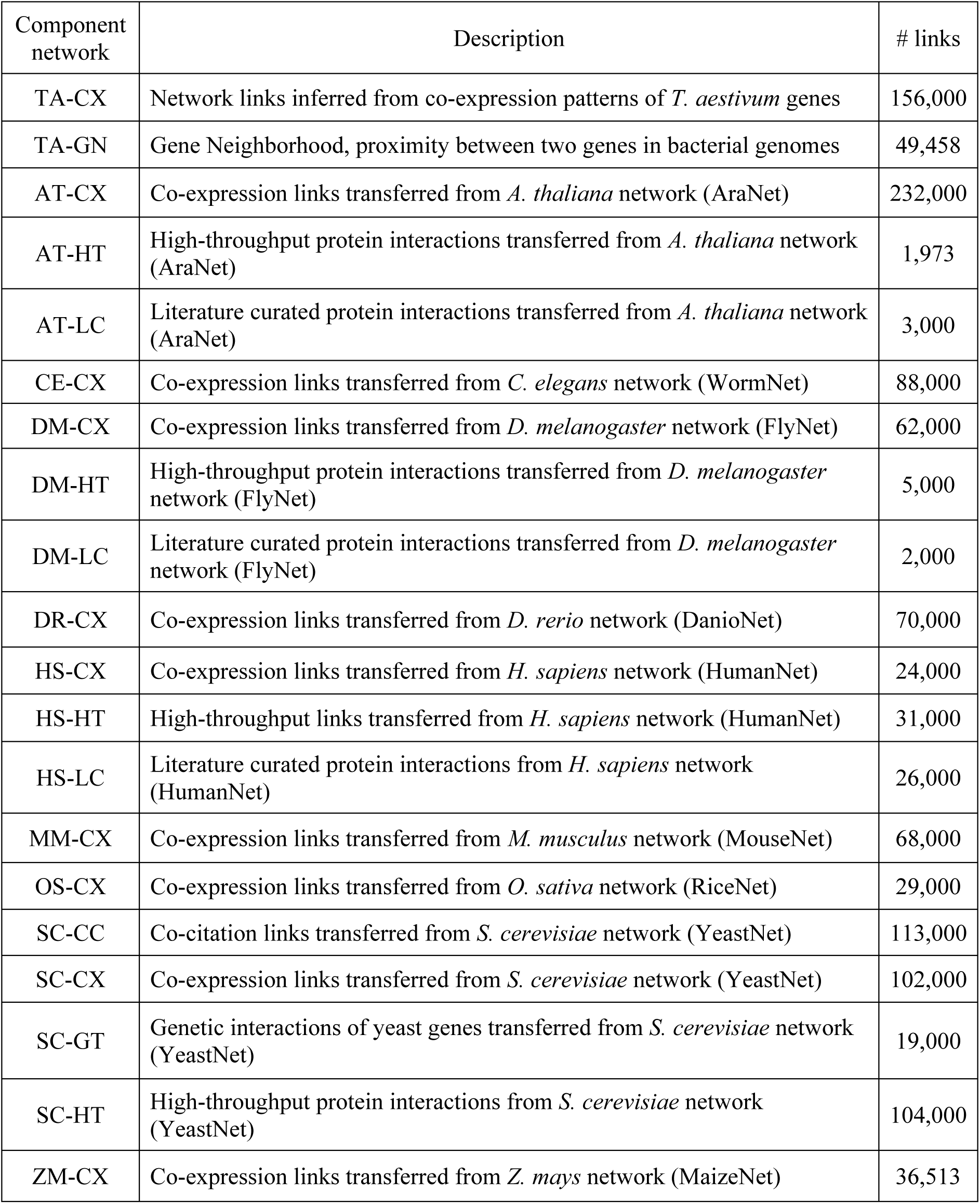
List of 20 component networks integrated to construct WheatNet

| Component network | Description | # links |
| --- | --- | --- |
| TA-CX | Network links inferred from co-expression patterns of *T. aestivum* genes | 156,000 |
| TA-GN | Gene Neighborhood, proximity between two genes in bacterial genomes | 49,458 |
| AT-CX | Co-expression links transferred from *A. thaliana* network (AraNet) | 232,000 |
| AT-HT | High-throughput protein interactions transferred from *A. thaliana* network (AraNet) | 1,973 |
| AT-LC | Literature curated protein interactions transferred from *A. thaliana* network (AraNet) | 3,000 |
| CE-CX | Co-expression links transferred from *C. elegans* network (WormNet) | 88,000 |
| DM-CX | Co-expression links transferred from *D. melanogaster* network (FlyNet) | 62,000 |
| DM-HT | High-throughput protein interactions transferred from *D. melanogaster* network (FlyNet) | 5,000 |
| DM-LC | Literature curated protein interactions transferred from *D. melanogaster* network (FlyNet) | 2,000 |
| DR-CX | Co-expression links transferred from *D. rerio* network (DanioNet) | 70,000 |
| HS-CX | Co-expression links transferred from *H. sapiens* network (HumanNet) | 24,000 |
| HS-HT | High-throughput links transferred from *H. sapiens* network (HumanNet) | 31,000 |
| HS-LC | Literature curated protein interactions from *H. sapiens* network (HumanNet) | 26,000 |
| MM-CX | Co-expression links transferred from *M. musculus* network (MouseNet) | 68,000 |
| OS-CX | Co-expression links transferred from *O. sativa* network (RiceNet) | 29,000 |
| SC-CC | Co-citation links transferred from *S. cerevisiae* network (YeastNet) | 113,000 |
| SC-CX | Co-expression links transferred from *S. cerevisiae* network (YeastNet) | 102,000 |
| SC-GT | Genetic interactions of yeast genes transferred from *S. cerevisiae* network (YeastNet) | 19,000 |
| SC-HT | High-throughput protein interactions from *S. cerevisiae* network (YeastNet) | 104,000 |
| ZM-CX | Co-expression links transferred from *Z. mays* network (MaizeNet) | 36,513 |

**Supplemental Table 2.** List of nine GEO series (GSE) used to infer co-expression links between wheat genes

| GEO Series | Description | # links |
| --- | --- | --- |
| GSE6027 | Meiosis and microsporogenesis microarray expression analysis | 71,000 |
| GSE6227 | Expression profiling after *Puccinia triticina* inoculation to resistant and susceptible lines | 86,000 |
| GSE9915 | Expression profiling after *Puccinia triticina* inoculation | 61,000 |
| GSE11774 | Microarray expression analysis of cold treated wheat cultivars | 97,000 |
| GSE23889 | Expression analysis of cold treatment | 105,000 |
| GSE31753 | Expression profiling after *Puccinia striiformis* f. sp. *Tritici* infection | 52,000 |
| GSE31756 | Expression profiling after *Puccinia striiformis* f. sp. *Tritici* infection | 84,000 |
| GSE54260 | Expression profiling of powdery mildew infection | 46,000 |
| GSE69437 | Expression profiling of *Fusarium graminearum* infection | 37,000 |

**Supplemental Figure 1.**
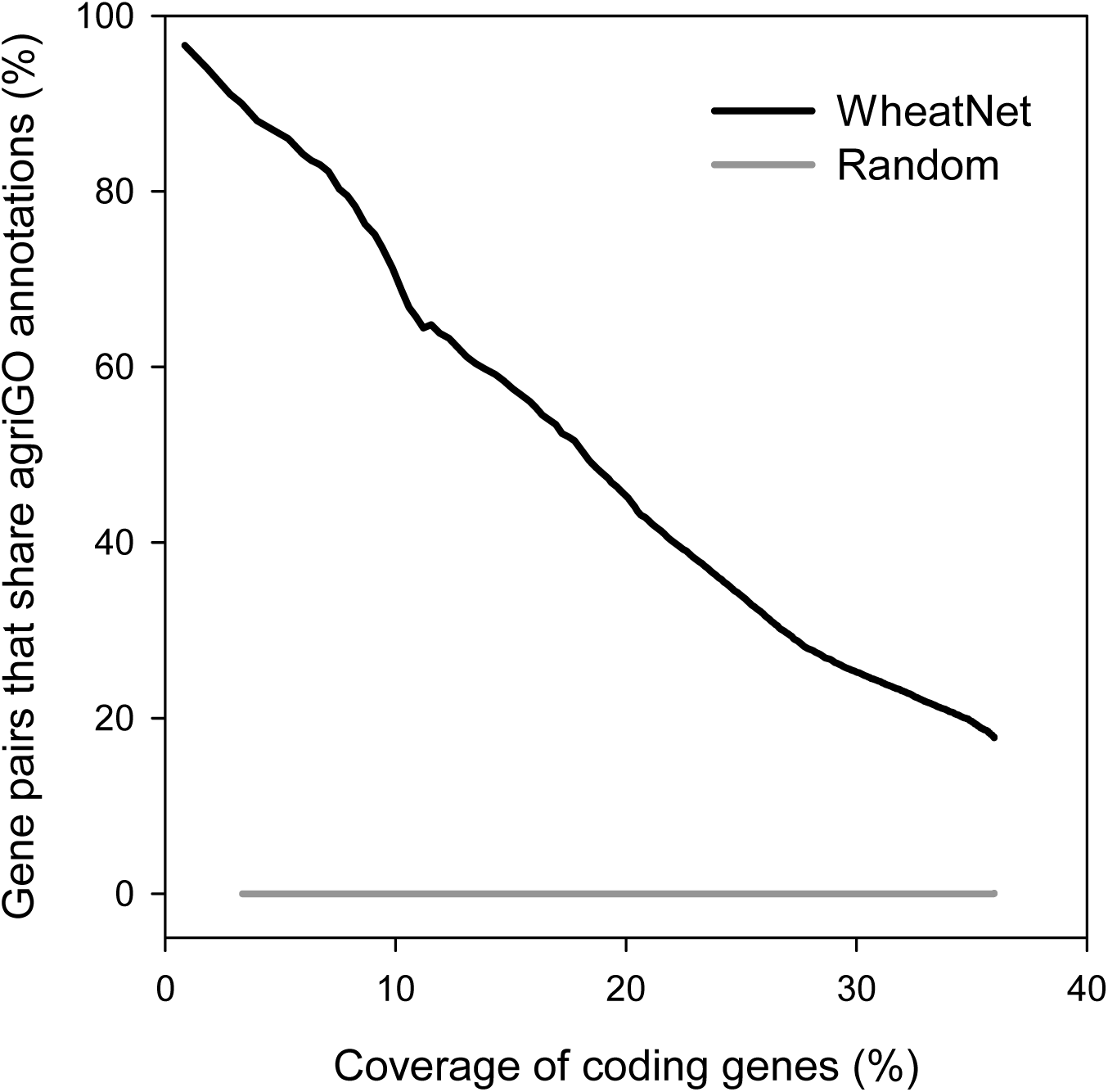
**Functional links of WheatNet are highly accurate.** To assess networks with validation data that are independent from those for network training, we used GOBP annotations by agriGO. Top-ranked links of WheatNet connect gene pairs of which >95% of them have at least one shared agriGO annotation between them, whereas only few of random gene pairs have shared annotations. The accuracy decreases as the network size increases.

## References

Cheng, Z., Dong, K., Ge, P., Bian, Y., Dong, L., Deng, X., Li, X., and Yan, Y. (2015). Identification of Leaf Proteins Differentially Accumulated between Wheat Cultivars Distinct in Their Levels of Drought Tolerance. PLoS One 10:e0125302.

Du, Z., Zhou, X., Ling, Y., Zhang, Z., and Su, Z. (2010). agriGO: a GO analysis toolkit for the agricultural community. Nucleic acids research 38:W64–70.

International Wheat Genome Sequencing, C. (2014). A chromosome-based draft sequence of the hexaploid bread wheat (Triticum aestivum) genome. Science 345:1251788.

Lee, I., Ambaru, B., Thakkar, P., Marcotte, E.M., and Rhee, S.Y. (2010). Rational association of genes with traits using a genome-scale gene network for Arabidopsis thaliana. Nat Biotechnol 28:149–156.

Liu, Z., Xin, M., Qin, J., Peng, H., Ni, Z., Yao, Y., and Sun, Q. (2015). Temporal transcriptome profiling reveals expression partitioning of homeologous genes contributing to heat and drought acclimation in wheat (Triticum aestivum L.). BMC Plant Biol 15:152.

Lopes, C.T., Franz, M., Kazi, F., Donaldson, S.L., Morris, Q., and Bader, G.D. (2010). Cytoscape Web: an interactive web-based network browser. Bioinformatics 26:2347–2348.

Mandal, M.S., Fu, Y., Zhang, S., and Ji, W. (2014). Proteomic analysis of the defense response of wheat to the powdery mildew fungus, Blumeria graminis f. sp. tritici. Protein J 33:513–524.

Uauy, C., Paraiso, F., Colasuonno, P., Tran, R.K., Tsai, H., Berardi, S., Comai, L., and Dubcovsky, J. (2009). A modified TILLING approach to detect induced mutations in tetraploid and hexaploid wheat. BMC Plant Biol 9:115.

Upadhyay, S.K., Kumar, J., Alok, A., and Tuli, R. (2013). RNA-guided genome editing for target gene mutations in wheat. G3 3:2233–2238.

Wojcik, P.I., Ouellet, T., Balcerzak, M., and Dzwinel, W. (2015). Identification of biomarker genes for resistance to a pathogen by a novel method for meta-analysis of single-channel microarray datasets. J Bioinform Comput Biol 13:1550013.

## Supplemental References

Barrett, T., Wilhite, S.E., Ledoux, P., Evangelista, C., Kim, I.F., Tomashevsky, M., Marshall, K.A., Phillippy, K.H., Sherman, P.M., Holko, M., et al. (2013). NCBI GEO: archive for functional genomics data sets--update. Nucleic acids research 41:D991–995.

Buchfink, B., Xie, C., and Huson, D.H. (2015). Fast and sensitive protein alignment using DIAMOND. Nat Methods 12:59–60.

Cho, A., Shin, J., Hwang, S., Kim, C., Shim, H., Kim, H., Kim, H., and Lee, I. (2014). WormNet v3: a network-assisted hypothesis-generating server for Caenorhabditis elegans. Nucleic acids research 42:W76–82.

Human Microbiome Project, C. (2012). Structure, function and diversity of the healthy human microbiome. Nature 486:207–214.

Kim, E., Hwang, S., Kim, H., Shim, H., Kang, B., Yang, S., Shim, J.H., Shin, S.Y., Marcotte, E.M., and Lee, I. (2016). MouseNet v2: a database of gene networks for studying the laboratory mouse and eight other model vertebrates. Nucleic acids research 44:D848–854.

Kim, E., Kim, H., and Lee, I. (2013). JiffyNet: a web-based instant protein network modeler for newly sequenced species. Nucleic acids research 41:W192–197.

Kim, H., Shin, J., Kim, E., Kim, H., Hwang, S., Shim, J.E., and Lee, I. (2014). YeastNet v3: a public database of data-specific and integrated functional gene networks for Saccharomyces cerevisiae. Nucleic acids research 42:D731–736.

Lee, I., Blom, U.M., Wang, P.I., Shim, J.E., and Marcotte, E.M. (2011). Prioritizing candidate disease genes by network-based boosting of genome-wide association data. Genome research 21:1109–1121.

Lee, I., Date, S.V., Adai, A.T., and Marcotte, E.M. (2004). A probabilistic functional network of yeast genes. Science 306:1555–1558.

Lee, T., Kim, H., and Lee, I. (2015a). Network-assisted crop systems genetics: network inference and integrative analysis. Curr Opin Plant Biol 24:61–70.

Lee, T., Oh, T., Yang, S., Shin, J., Hwang, S., Kim, C.Y., Kim, H., Shim, H., Shim, J.E., Ronald, P.C., et al. (2015b). RiceNet v2: an improved network prioritization server for rice genes. Nucleic acids research 43:W122–127.

Lee, T., Yang, S., Kim, E., Ko, Y., Hwang, S., Shin, J., Shim, J.E., Shim, H., Kim, H., Kim, C., et al. (2015c). AraNet v2: an improved database of co-functional gene networks for the study of Arabidopsis thaliana and 27 other nonmodel plant species. Nucleic acids research 43:D996–1002.

Lehner, B., and Lee, I. (2008). Network-guided genetic screening: building, testing and using gene networks to predict gene function. Brief Funct Genomic Proteomic 7:217–227.

Li, L., Stoeckert, Jr. C.J., and Roos, D.S. (2003). OrthoMCL: identification of ortholog groups for eukaryotic genomes. Genome research 13:2178–2189.

O'Brien, K.P., Remm, M., and Sonnhammer, E.L. (2005). Inparanoid: a comprehensive database of eukaryotic orthologs. Nucleic acids research 33:D476–480.

Overbeek, R., Fonstein, M., D'Souza, M., Pusch, G.D., and Maltsev, N. (1999). The use of gene clusters to infer functional coupling. Proc Natl Acad Sci U S A 96:2896–2901.

Shim, H., Kim, J.H., Kim, C.Y., Hwang, S., Kim, H., Yang, S., Lee, J.E., and Lee, I. (2016). Function-driven discovery of disease genes in zebrafish using an integrated genomics big data resource. Nucleic acids research 44:9611–9623.

Shin, J., Lee, T., Kim, H., and Lee, I. (2014). Complementarity between distance- and probability-based methods of gene neighbourhood identification for pathway reconstruction. Mol Biosyst 10:24–29.

Shin, J., Yang, S., Kim, E., Kim, C.Y., Shim, H., Cho, A., Kim, H., Hwang, S., Shim, J.E., and Lee, I. (2015). FlyNet: a versatile network prioritization server for the Drosophila community. Nucleic acids research 43:W91–97.

Sunagawa, S., Coelho, L.P., Chaffron, S., Kultima, J.R., Labadie, K., Salazar, G., Djahanschiri, B., Zeller, G., Mende, D.R., Alberti, A., et al. (2015). Ocean plankton. Structure and function of the global ocean microbiome. Science 348:1261359.

Tello-Ruiz, M.K., Stein, J., Wei, S., Preece, J., Olson, A., Naithani, S., Amarasinghe, V., Dharmawardhana, P., Jiao, Y., Mulvaney, J., et al. (2016). Gramene 2016: comparative plant genomics and pathway resources. Nucleic acids research 44:D1133–1140.

Walhout, A.J., Sordella, R., Lu, X., Hartley, J.L., Temple, G.F., Brasch, M.A., Thierry-Mieg, N., and Vidal, M. (2000). Protein interaction mapping in C. elegans using proteins involved in vulval development. Science 287:116–122.

